# Maximizing vegetation representation of the catchment in sedimentary DNA with targeted cores in small lakes

**DOI:** 10.1101/2025.07.02.662717

**Authors:** Tulug Gulce Ataman, Youri Lammers, Inger Greve Alsos, Dilli Prasad Rijal, Antony G. Brown

## Abstract

Sedimentary DNA is becoming an invaluable tool for biodiversity assessments across spatial and temporal scales. Accurate interpretation, however, requires a clear understanding of its origin and taphonomy, from transport to preservation processes within lake systems. Insights into these processes are crucial for developing efficient sampling methods for precise ecological monitoring. Our analysis of 42 adjacent surface sediment sample replicates shows that deeper, central samples, with reduced influence from aquatic taxa, exhibit greater taxonomic richness compared to near-shore samples. By comparing these results to systematic vegetation surveys, we demonstrate that central cores are optimal as they capture the main taxonomic richness in the catchment, while marginal samples near inflows are essential for detecting rarer, spatially restricted taxa. This study highlights the potential of central-core sampling to effectively capture vegetation profiles in small, bathymetrically simple lakes, while enhancing the understanding of DNA transport mechanisms in catchments with similar topographic or hydrological characteristics.

## Introduction

The detection of sedimentary DNA (sedDNA) in lake sediments is a promising method in different fields of environmental study such as effect of climate and human land-use on vegetation ^1,2^, monitoring of rare or invasive species ^3,4^, and reconstructing past environmental conditions ^5,6,7^. Furthermore, sedimentary ancient DNA (*sed*aDNA), which is deposited in chronostratigraphic order in lake sediments, can be used to interpret changes in past ecosystems and biodiversity over time ^8^. However, these studies make a fundamental assumption that a single sampling location is representative of both the lake and the catchment. This appears to be in contradiction to some evidence from samples near archaeological sites that suggest that there can be local effects of human activity which decrease further out into the lake^2^. A recent study ^9^ using multiple lake-sediment cores from the large lake Constance reveals significant spatial variation in sedDNA distribution at the level of Molecular Operational Taxonomic units (MOTUs), influenced by rare DNA detectability, sediment heterogeneity and geographic variability in the lake’s ecosystem. However, testing the representativity of plant DNA requires an extensive vegetation survey and multiple sediment samples to provide high-resolution data for accurate assessment.

DNA may originate from living and decaying organisms and may be either transported into the lake or produced *in situ* ^10,11^. One of the challenges regarding the environmental DNA (eDNA) analysis of biodiversity is related to the limited knowledge of DNA taphonomy across ecosystems ^12,13^. Taphonomical concerns about DNA include depositional mechanisms of environmental DNA in sediments affected by geochemical processes ^14^ and post-mortem degradation/decomposition and transportation characteristics from its source to its deposition in sediments or any environmental samples ^15–17^. Rivers transport eDNA over distances that varies with stream velocity and dissolved oxygen concentrations ^18–20^, and it can be transported by being attached to small particles (clastic sediments or organic particles) and especially clay minerals, or in a free state, but this undergoes far higher rates of degradation ^21,22^.

Alsos *et al.* ^23^ compared vegetation surveys with a sediment samples taken from the central, deepest part of the lake and demonstrated that approximately 73% of plant species found in lake surface sediments matched those identified in botanical surveys within 2 m from the lake, with a further 12% matching records within 50 m from the lake shore. However, Alsos *et al.* ^23^ did not perform extensive vegetation surveys beyond 2 m, leaving uncertainties regarding the significance of the source (plant) distance from the lakes. Furthermore, a modified Kajak corer was used to collect two sediment cores per sampling point and the difference between these samples was found to be insignificant, but further within-lake variation was not explored. Similarly, Wang *et al.*^9^ reported that the transport mechanisms of plant sedDNA in the lake involved both water-mediated pathways and localized deposition. They noted that common plants exhibited widespread distribution, while rarer taxa displayed specific distribution patterns near the lake’s inflow, but lacked detailed vegetation surveys to further explore these patterns. These findings collectively underscore the partial understanding of the spatial variation of sedDNA within a lake suggesting that a systematic sampling approach is needed to better understand spatial variation of sedDNA.

Here we investigate spatial sedDNA variation in a small lake, comparing it to detailed vegetation surveys from a catchment with diverse topography and vegetation types. We aim to understand the spatial patterns of sampled sedDNA and its relationship to surrounding vegetation by systematically sampling surface sediments across the lake and conducting detailed vegetation surveys in the catchment. More specifically, we address four research questions: **1)** Is there consistency between sampling point replicates, **2)** How many samples are needed to capture the majority of the taxa in the vegetation **3)** Does the abundance of aquatic vascular plants in sedDNA samples reduce detection of terrestrial plants, **4)** How does the spatial detection of DNA in the lake relate to the vegetation within the catchment?

Our results demonstrate that sedDNA in small lakes reflects catchment vegetation patterns with both consistency and spatial variation. Replicate samples taken within close proximity (0.15 m) yield similar taxonomic profiles, while larger-scale sampling (25–35 m) reveals systematic variation driven by catchment vegetation distribution and topography. Samples from depocenters exhibit the highest richness, particularly of terrestrial taxa, whereas areas dominated by aquatic plants near the shore suppress terrestrial signals. Our findings establish central-core sampling as a reliable method for documenting dominant vegetation profiles in small catchments with simple lake bathymetry. This study demonstrates how elucidating DNA transport mechanisms enables biodiversity monitoring to efficiently capture vegetation diversity with reduced field efforts.

## Results

### Molecular data

Stabbevatnet, a small lake in northern Norway (Figure 1), was selected for its contrasting vegetation zones and sampled along 12 transects covering fluvial deposition pathways of DNA. 84 surface sediment samples were systematically collected across 12 transects with two replicates at each of the 42 sampling points (Figure 1). After DNA extraction of the samples and controls, amplification of the p6-loop region and sequencing, we obtained a total of 42,463,283 raw reads. After identification and filtering, this was reduced to 15,361,680 reads belonging to 196 barcodes with a 100% sequence similarity to at least one of the reference libraries (Supplementary Data 1). Taxonomic assignments were manually inspected and categorized, and only high-confidence taxa were retained for further analysis. Of these sequences, 154 were assigned to category 1 (assumed true positive), 41 to category 2 (likely true positive), and 27 to category 3 (assumed false positive). Sequences in categories 1 and 2 were retained and those assigned to the same taxa were collapsed resulting in a final set of 165 taxa (Supplementary Data 1).

**Figure 1.**
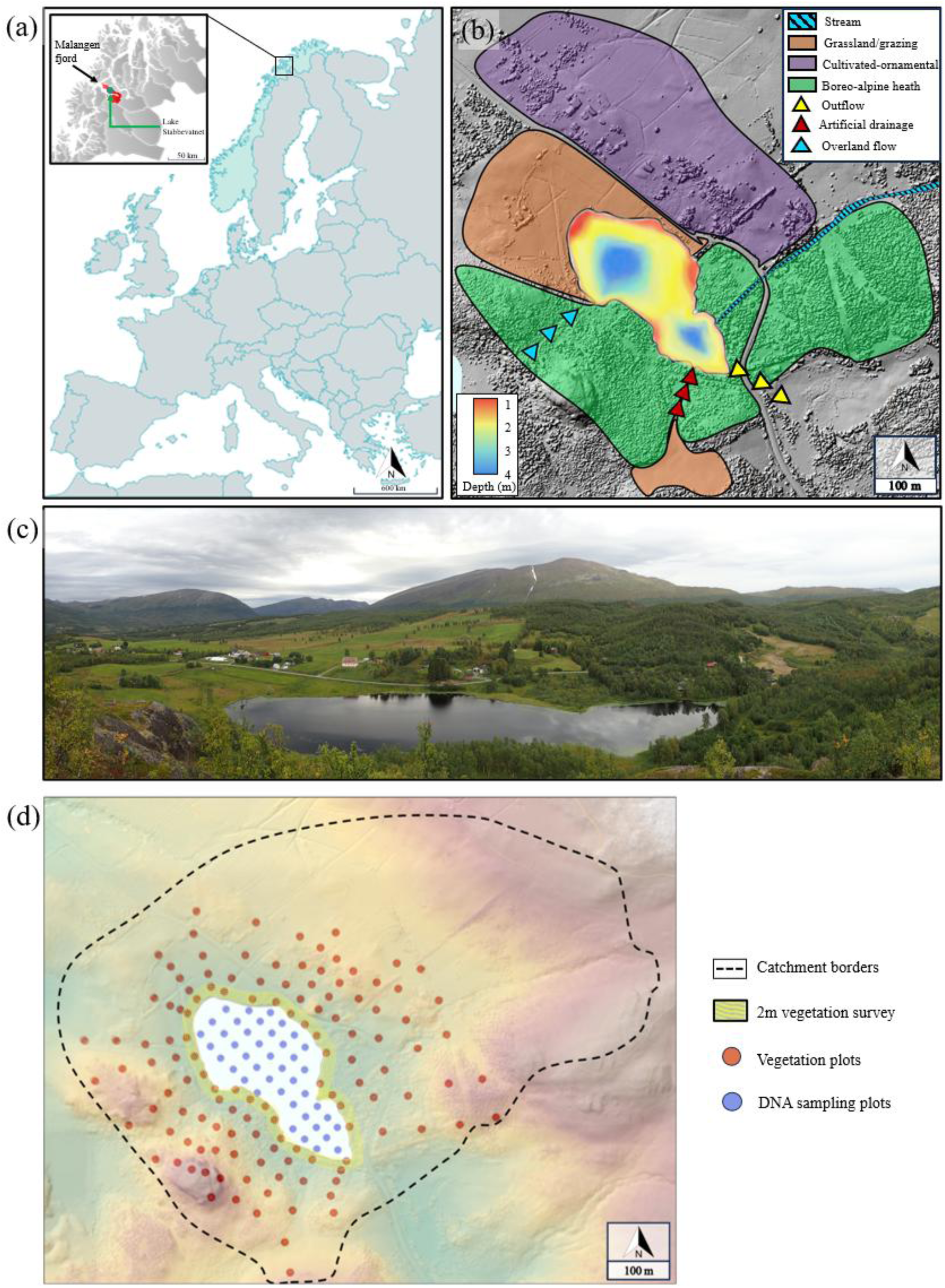
Study site and vegetation profile. **(a)** Location of the study site Stabbevatnet, county of Troms, northern Norway. **(b)** The vegetation map around the lake orange; Grassland or grazing vegetation, purple; Cultivated plants and ornamental species, green; Boreo-alpine heath with a rich willow (*Salix* sp.), spruce (*Picea abies*) and birch (*Betula nana, B. pubescens*) forest. **(c)** Stabbevatnet lake viewed from the south-west. Photo: Marie Føreid Merkel. **(d)** Systematic sampling from sediment, vegetation plots within the catchment borders and 2 m vegetation survey.

### Comparison of sample replicates and richness

We detected an average of 55.8 ± 19.7 and 56.5 ± 21.0 taxa per sample in replicate datasets a and b, respectively (Supplementary Data 2, columns *richness_a* and *richness_b*). Based on the taxonomic assignments, the average Jaccard similarity between the sample replicates was 59% (Supplementary Figure 1; Supplementary Data 2), with only 10 replicates having a similarity below 50% (Supplementary Data 2). We therefore merged the taxa detected in the replicate samples resulting in a total of 115 taxa that were detected in both samples simultaneously (Supplementary Data 3). The most abundant taxa across the 42 DNA samples, such as *Betula* and *Filipendula ulmaria*, were consistently present in both replicates (Supplementary Data 3).

A breakpoint analysis was carried out on the taxa accumulation curves for the 42 merged DNA samples to estimate the optimal sampling number to capture the diversity. The breakpoint in the accumulation curve indicates only a moderate increase in taxa after sampling 8.3 samples (Figure 2a). At this breakpoint, we observed an average richness of approximately 82.34 (±9.46) taxa, which is 73% of the total taxa present in the merged data (Supplementary Data 3). The average richness breakpoint is comparable to the highest richness observed in a single sample which is 82 taxa (Figure 2). The highest richness observed in a single merged sample was 82 taxa (Supplementary Data 3).

**Figure 2.**
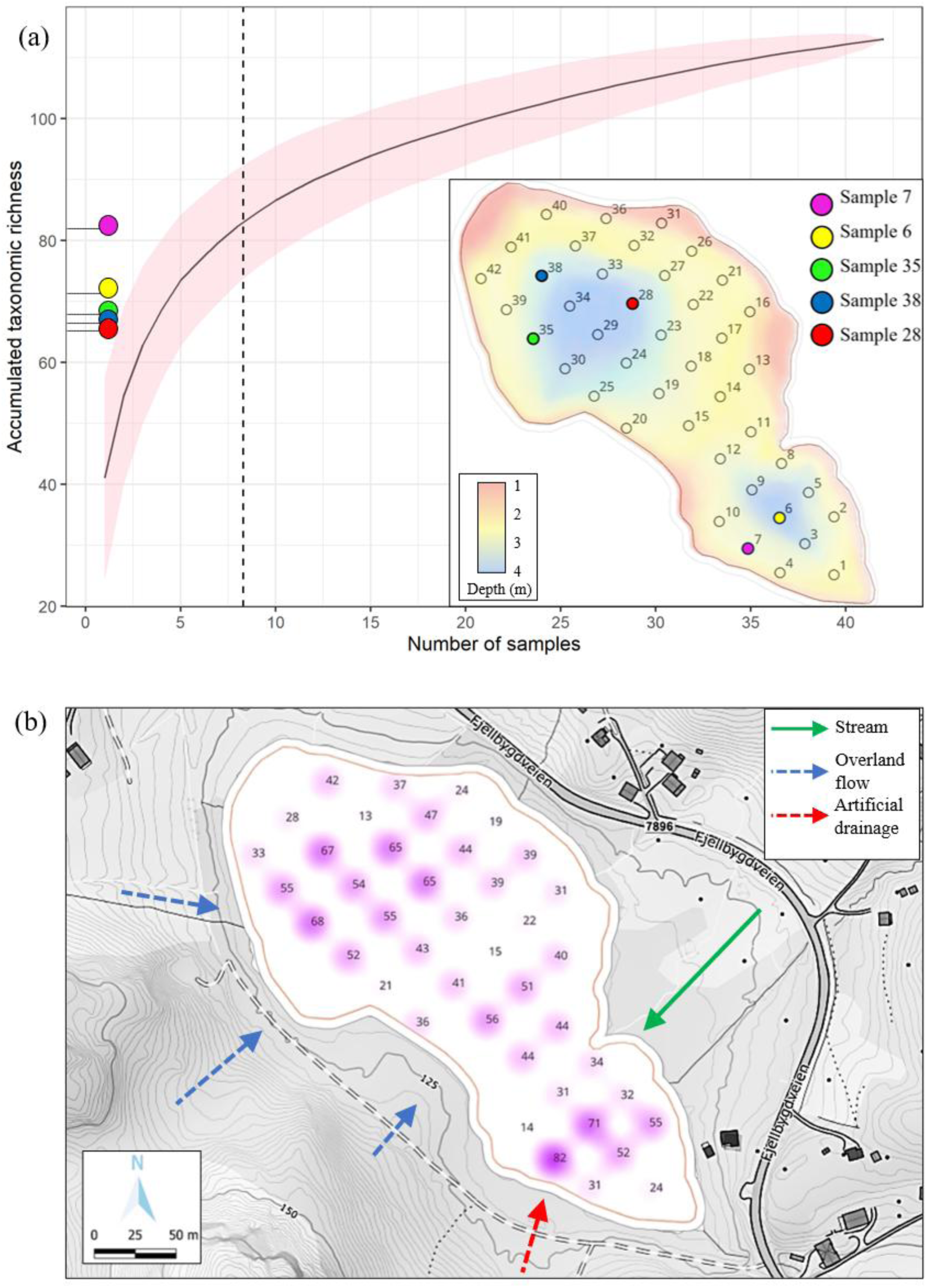
Accumulated Taxonomic Richness and Shared Taxa Patterns in Relation to Sediment Samples and Fluvial Inputs. **(a)** Mean accumulated richness curve based on 1000 randomization. The vertical dashed line indicates the number of sediment samples required to capture the optimal number of taxa. The five samples with the highest richness are indicated on the y-axis. The insert map shows the location of the five samples with the highest richness. **(b)** Richness based on shared taxa in A and B samples and the fluvial inputs to the lake.

### Spatial distribution of richness

The taxonomic richness ranged from 14 to 82 taxa in the merged samples with an average of 40.95 (±16.21) taxa. A higher richness was observed in the samples located from the deepest parts of the lake and/or near the inflow streams (Figure 2b). The highest richness of 82 was observed in sample 7, located near the southwestern artificial drainage inflow (Figure 2). The samples located in the shallower parts of the lake had both a lower total richness (terrestrial taxa richness and aquatic taxa richness), as well as a higher proportion of aquatic reads (Figure 2b; Figure 3a). Taxa such as *Myriophyllum sibiricum* and *Nuphar pumila* were observed in samples with up to 97% and 86% of the reads respectively (Figure 3b, Supplementary Data 4). We observed a statistically significant negative correlation between the proportion of aquatic reads and both total richness (r = -0.45, p-value < 0.01) and proportion of terrestrial taxa richness (r = - 0.69, p-value < 0.0001), suggesting that an increase in the proportion of aquatic reads is associated with reduced taxonomic diversity, particularly among terrestrial taxa (Figure 3; Supplementary Figure 2).

**Figure 3.**
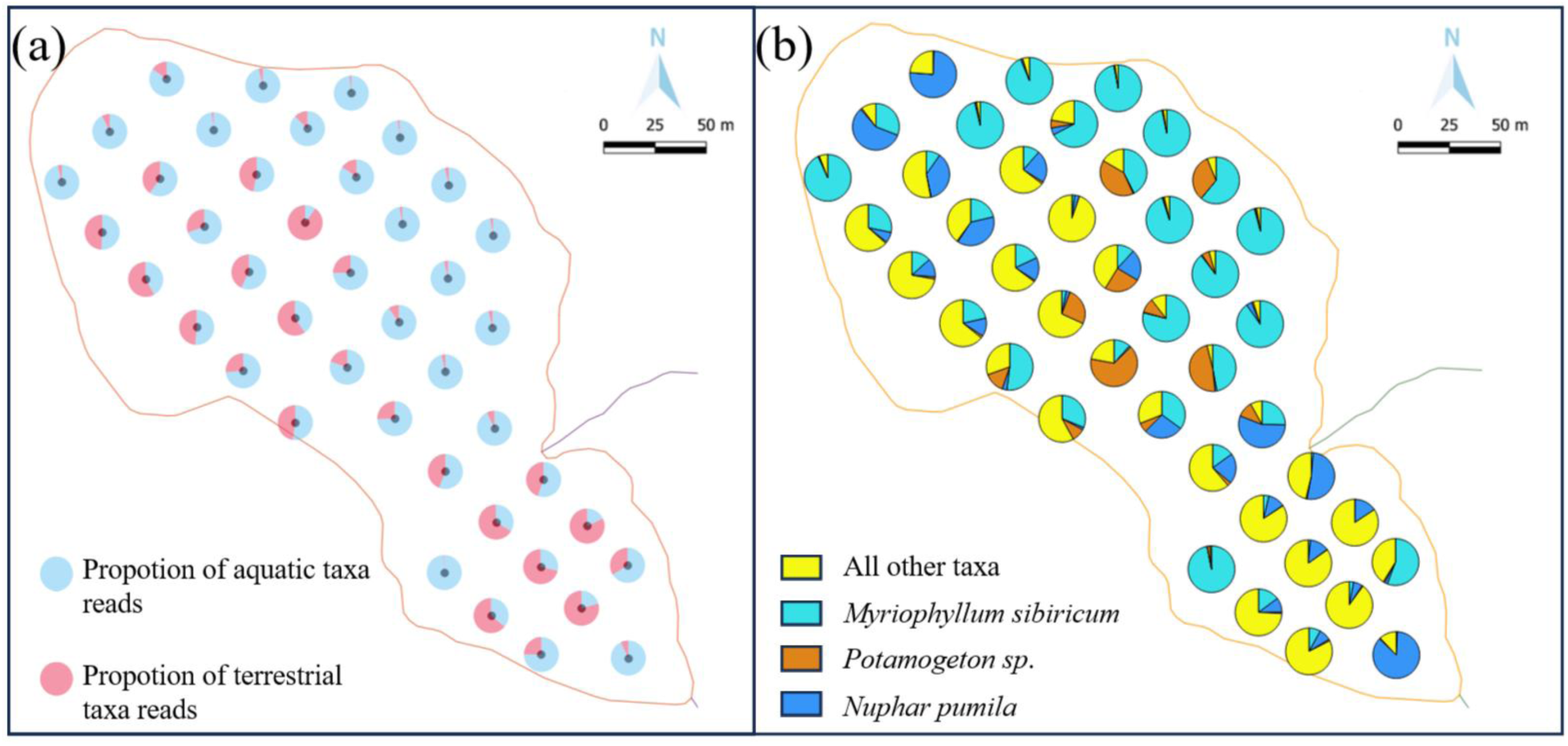
Proportion of aquatic and terrestrial plant reads based on 42 merged DNA samples. **(a)** Each pie chart displays the relative read proportions of the aquatic (blue) and terrestrial (red) taxa across the sampled locations. **(b)** Each pie chart displays the relative read proportions for the three most abundant aquatic taxa and all other taxa combined, across the sampled locations.

A significant negative correlation was observed between the proportion of aquatic reads and replicate similarity across sediment samples (Pearson’s r = -0.54, p < 0.001; Supplementary Figure 3). Linear regression analysis confirmed a decline in replicate similarity with increasing aquatic read proportion (y = -0.24x + 0.75, R² = 0.29).

### Vegetation surveys

The vegetation in a total of 129 1 m² vegetation plots across the lake’s catchment was surveyed. Across all vegetation plots, 139 taxa were recorded, yielding 1119 total observations (Supplementary Data 4). The average number of taxa per vegetation plot was 8.7 ± 5.3 (Supplementary Data 4). *Betula* was the most frequently detected taxon, observed in 61 vegetation plots.

A second survey within 2 m of the lakeshore, targeting taxa likely contributing most to sedDNA, classified taxa abundance from rare to dominant^23^ (Figure 4a). Additional taxa observations were recorded within the catchment, beyond the 1 m² vegetation plots and the 2 m vegetation survey range (Supplementary Data 5). In the 2 m vegetation survey, 84 taxa were recorded with an average abundance value of 2.3. Combined across both vegetation surveys a total of 155 taxa were recorded and 110 specimens were collected for identification and preservation in the Tromsø University Museum herbarium.

**Figure 4.**
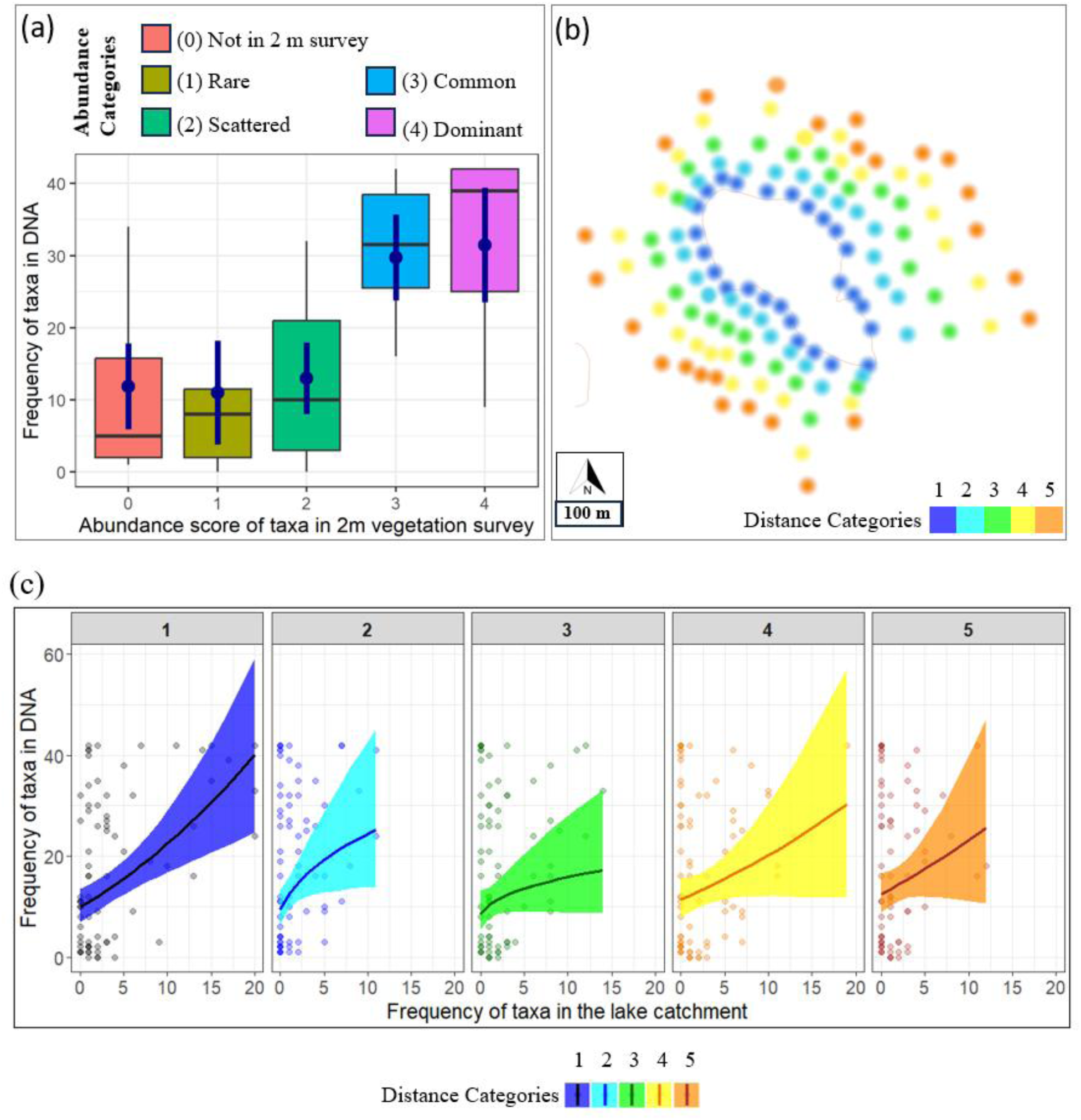
Relationship between frequency (number of detections) of taxa in the catchment to frequency based on detection in the sedDNA samples. **(a)** Association between DNA frequencies and abundance of taxa recorded in a 2 m vegetation survey**. (b)** Distance categories of vegetation plots based on their distance to the lake **(c)** Associations between vegetation frequencies from different distance categories (shown in **b**) of the catchments (x-axis) and DNA frequencies from the lake sediment (y-axis). The lines and shadings represent fitted values ± 2SE in their original scales.

### Comparison with the vegetation data

To align sedDNA results with vegetation data, both lists were standardized to the highest taxonomic level observed. After species list standardization with DNA data (Supplementary Data 1), the number of taxa recorded in the vegetation surveys was condensed to 89 unique taxa (Supplementary Data 4). Out of the 89 standardized taxa in the sedDNA results (Supplementary Data 4; Supplementary Data 1), 18 were removed because they were not replicated between the A and B samples, resulting in a final set of 71 taxa in sedDNA data matching the vegetation taxa. To enable comparison, sedimentary DNA (sedDNA) frequency was defined as the number of sediment samples (out of 42) in which each taxon was detected. Vegetation frequencies were similarly calculated as the number of plots within each distance category where the same taxon was recorded. The frequencies from both surveys (the 2 m vegetation survey abundance and vegetation plots) were compared with sedDNA frequencies, taking vegetation plot distance to the lake into account. The frequency of taxa in DNA showed a positive association with their frequency in the catchment vegetation across various distances from the lakeshore. A positive association was observed between sedDNA frequency and higher abundance categories (i.e. common and dominant) from the 2 m vegetation survey where over 31% of the variation in the DNA frequency was explained by vegetation frequency (F_4,70_ = 9.39, R^2^ = 0.31, p < 0.0001, Supplementary Table 1). The positive association between DNA detection and catchment vegetation tended to decrease with distance from the lakeshore. Vegetation frequency in the nearest zone (Distance Category 1), located approximately 0 to 3 m from the lakeshore, explained 16% of the variation in DNA frequency representing the highest association among all distance categories (F_1,73_ = 15.1, R^2^_adj_ = 0.16, p < 0.001, Supplementary Table 2). The highest overall association was observed when vegetation data from all distance categories were combined (F_1,84_ = 26.48, R^2^_adj_ = 0.23, p < 0.0001, Supplementary Table 2).

## Discussion

The 42 merged DNA samples from Lake Stabbevatnet indicate a high similarity between the samples. The spatial distribution of these similarity values reveal that the greatest similarity is observed in the two depocenters of the lake (Supplementary Figure 1). The replicates with the lowest similarity generally occurred in areas closer to the shore (Supplementary Figure 1b). Although this similarity may be affected by the domination of aquatics (Supplementary Figure 3) in some samples or technical issues such as a PCR replicate that failed or low raw read counts in other samples ^24,25^. The high similarity observed between surface replicates supports the method of creating compound sequences out of parallel sediment cores.

The pattern of total richness varies systematically across the lake, with the highest values detected in the two depocenters and on the steeper southwest side of the lake. This may be caused by a movement of sediment from the major input streams across the lake towards the outlet and possibly due to a wave effect as the topographic asymmetry of the basin favors cross-lake winds from the north and northwest to the south and southeast rather than winds from the southwest. The highest richness location, with over 72% of the total richness, was near to a drainage outlet that drained both agricultural grassland, woodland and shrubland and had the highest slope close to the lake (Figure 2b). The area with the lowest richness is the northeastern shallow shelf area, possibly caused by a lack of sediment accumulation. The overall taxa richness profile showed a general increase on the western side of the lake compared to the eastern side, with the highest richness observed near steep slopes and major drainage outlets. This pattern could be influenced by erosion processes and topographic features, such as steep slopes and well-developed drainage networks, which contribute to terrestrial taxa richness in sediment DNA through the transport of adsorbed DNA ^16^. Therefore, a high-resolution spatial analysis that also includes elevation alongside longitude, and latitude data for each vegetation plot and comparing the occurrence of taxa in these plots vs in sedimentary DNA may give an additional insight for DNA transportation pathways.

The breakpoint analysis indicates that eight random samples are required to capture the majority of the taxa detected in the lake. However, a single sample, located in a high richness area, such as the depocenters or inflow locations, recovers a similar number of taxa. Dominant taxa in the vegetation are detected in the majority of the DNA samples, regardless of their location in the lake (Figure 5a). Rare taxa on the other hand, can have a more restricted distribution in the lake (Figure 5b). Thus, a single sample from a high richness area captures the common and abundant taxa but might not be able to recover the rarer taxa. If the aim is to obtain a full record of all taxa in the catchment, multiple samples from different locations are required. Spatial variation and localization of taxa should be considered during sampling and analysis, as deeper cores in one basin are likely to have a better chance of capturing the rare taxa of vegetation near that basin.

**Figure 5.**
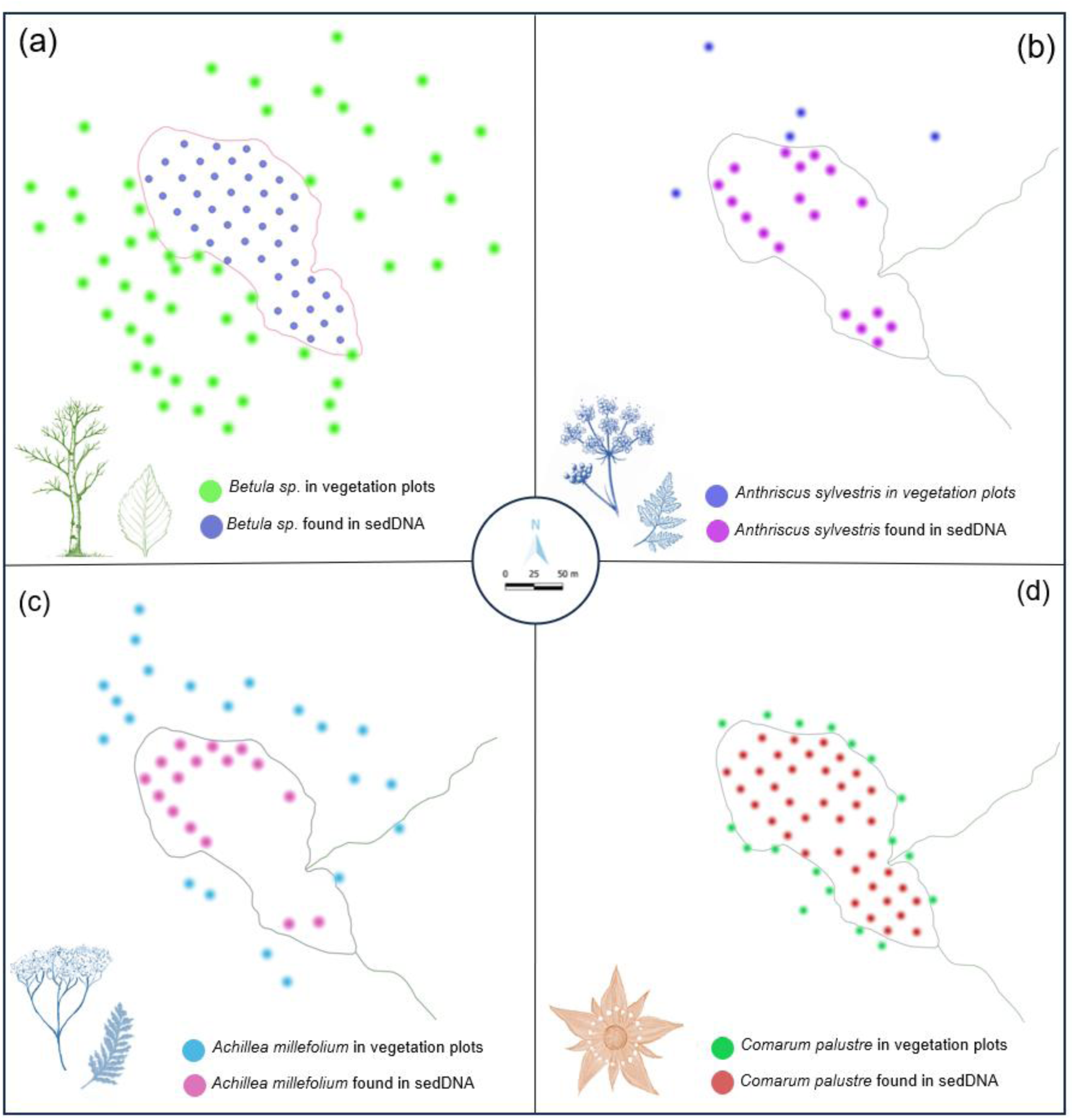
Spatial distribution of taxa found in DNA and vegetation plots. **(a)** *Betula sp.* found in vegetation plots and sedDNA **(b)** *Anthriscus sylvestris* found in vegetation plots and in sedDNA **(c)** *Achillea millefolium* found in vegetation plots and in sedDNA **(d)** *Comarum palustre* found in vegetation plots and in sedDNA

The presence of aquatic taxa can negatively impact the detection of terrestrial plants and total richness (Figure 3, Supplementary Figure 3). This is possibly due to aquatic plant remains being present in the sediments and subsequent DNA extraction. The high concentration of aquatic DNA is likely to outcompete the less abundant terrestrial DNA during PCR amplification, resulting in an underrepresentation of terrestrial plant richness. Taxa such as *Myriophyllum sibiricum*, *Nuphar pumila* and *Potamogeton* could be detected with >90% of the reads. These taxa were generally found in shallower, yet distinct, areas of the lake. *M. sibiricum* was primarily found along the northern edge of the northern depocenter, *N. pumila* was detected near the in- and outflow of the lake, along with the north-western side of the lake. Finally, *Potamogeton* was detected in the shallow area between the two depocenters. In order to avoid aquatic DNA swamping, one should core in locations with less aquatic plant growth, such as deeper parts of a lake.

The probability of detecting terrestrial plant taxa in DNA samples varied across the lake, influenced by the DNA transportation patterns and the distribution of plant taxa in the catchment (Figure 4; Figure 5). Common taxa in the vegetation plot survey, such as willow (*Salix*) and birch (*Betula*; Figure 5a), are detected in all sedDNA samples. This is also true for abundant taxa in the 2 m vegetation survey, or those abundantly found in the vegetation plots closest to the lake (Figure 4, Supplementary Data 4), such as meadowsweet (*Filipendula ulmaria*), *Caltha palustris and Comarum palustre* (Figure 5b) although they were not abundant in other distance categories further in the lake. The zoning of the terrestrial vegetation data shows that there is a clear association with distance from the lake and probability of detection in the lake. Overall, abundant taxa within the 2 m of the lakeshore are more likely to be detected compared to the rarer taxa. These findings align with those of Alsos et al. (2018) ^23^, who reported correlations between taxa detected from sedimentary DNA and vegetation within 2-10 m of the lakeshore, and also support assumptions made by Giguet-Covex et al. (2019) ^16^. The plants with a more restricted distribution around the lake were found in fewer sedDNA samples and some had a pattern of proximal association, for example yarrow (*Achillea millefolium*) (Figure 5c) and cow parsley (*Anthyriscus sylvestris*) (Figure 5d). These associations along with the distribution of richness suggests that there are preferred transport pathways for sedDNA through the stream and drainage network of the catchment ^16,26^.

Our study of Lake Stabbevatnet supports findings from Lake Constance^9^, showing that fluvial inputs play a significant role in transporting DNA into lake sediments and shaping localized sedimentary DNA patterns. While fluvial transport appears to contribute to sedimentary DNA regardless of catchment size, the catchment’s size and topography, together with the structure of the lake basin, may influence how DNA moves through the system and how far it travels before deposition. These factors can affect both the quantity and the composition of DNA preserved in the sediment. For example, in lakes with larger catchments, such as Lake Constance^9^, DNA is likely transported over longer distances, potentially integrating plant DNA from extensive surrounding areas across Austria, Germany, and Switzerland.

## Conclusion

We show here that closely spaced surface sediment sample replicates (0.15 m apart) have overall statistically indistinguishable sedDNA diversity values. However, replicate similarity decreases with increasing dominance of aquatic taxa, suggesting that spatial variability in sedDNA composition is greater in samples with higher aquatic input. Since samples from shallow, near-shore areas are dominated by aquatic taxa, the broader differences observed among samples spaced 25-35 m apart likely reflect local environmental variation, such as differences in depth and aquatic plant dominance. The sediment samples from the two lake depocenters had the highest richness. To improve the detection of terrestrial taxa and minimize the impact of aquatic plant DNA, it is beneficial to take cores from areas with less aquatic growth, such as the deeper parts of a lake.

Based on these results and the previous studies relating vegetation to sedimentary DNA data, we can broadly support the single-core approach to *sed*aDNA-based palaeoecological studies of small lakes. We also show that the lake’s DNA assemblage best matches the vegetation from < 2 m of the lake with declining representation of more distant taxa. However, spatial patterns of sedimentary DNA may vary in relation to distance to the shore, lake geography, water depth, catchment topography, and land use patterns within the lake catchment. Where there is a spatial pattern related to fluvial inputs, targeted cores near the fluvial input may provide additional ecological data. In summary, while deeper lake cores may reveal higher terrestrial taxa richness, the analysis of multiple cores will increase the chance of detecting rare taxa in the catchment and spatially restricted land use patterns.

## Methods

### Study site and Sampling Methods

Stabbevatnet (69°23’15.5“N,18°42’32.0”E) lies at 123 m asl (Norgeskart) in Malangen municipality in northern Norway (Figure 1). The lake has a surface area of approximately 0.043 km^2^ and has two basins, one in the northern part of the lake with a maximum water depth of 4 m and a second in the southern part of the lake with a maximum water depth of 5 m. The lake was chosen on the following criteria: **1)** the size and depth are comparable to many lakes used in sedimentary ancient DNA (sedaDNA) studies, **2)** the pattern of spatial variation both around the lake and in the catchment facilitates comparison with vegetation sedDNA detection in the lake. This is because the lake has arable and grazing fields to the north and east of the lake and boreo-alpine unenclosed heath with mixed forest (*Salix, Picea*, and *Betula*) on the west and southern sides of the lake (Figure 1). The lake has four primary inflows from the west and northwest (overland flow) and northeast (stream), and southwest (artificial drainage) and one outflow in the southeastern part of the southern basin (Figure 1a). The bathymetry map was created during sampling using a Helix 5 Chirp GPS G2 Digital Sonar (Hummingbird technologies, UK).

Catchment boundaries were initially delineated using NEVINA (http://nevina.nve.no) with a 10 m digital elevation model, testing multiple pour-point placements to evaluate sensitivity to local topography (Supplementary Materials; Supplementary Figure 4). However, the resulting catchments varied considerably in size and shape and often included areas beyond the lake’s local depositional zone (Supplementary Figure 4a). We therefore manually defined the catchment based on visible topographic features such as ridgelines and slope direction, supported by field observations and the spatial extent of vegetation plots (Supplementary Figure 4b). The final catchment (0.6 km²) includes the local basin shaped by natural elevation gradients, representing the area most likely to contribute terrestrial DNA to the lake and where vegetation composition aligns with both field surveys and sedimentary DNA. The final boundary (0.6 km²) reflects the locally enclosed drainage area that is consistent with ecological relevance of source of sedimentary DNA (Supplementary Figure 4b; Figure 1c). Expanding the catchment beyond this would include ecologically irrelevant terrain and could misrepresent local vegetation influence as taxa beyond this boundary were already represented within the defined catchment area (Supplementary Figure 4b; Figure 1c), and no new taxa were detected further upslope. Detailed justifications and comparison maps are provided in the Supplementary Materials (Supplementary Figure 4).

Stabbevatnet is situated in the Caledonian nappes region of Northern Norway (https://www.ngu.no). The bedrock is part of the Tromsø Nappe Complex and predominantly granitic muscovite-rich gneiss but covered by variable thicknesses of morainic sediments. The northern and eastern slopes of the catchment are covered by thick morainic superficial sediments which allows farming including cultivation despite the area being well above the maximum marine limit (60 m asl) which normally limits agriculture in N Norway. The defined catchment (0.6 km²) receives a mean annual precipitation of 1305 mm and has a mean annual temperature of 1.5 °C, with a mean July temperature of 11 °C, based on 1991-2020 climate normals calculated by NEVINA for this area. Land cover within the catchment consists of 44.4% forest and 41.6% cultivated land. Elevation ranges from 123 to 206 m, with an average slope of 8.7°, and the drainage density is 1.6 km^-1^, indicating moderately dissected terrain (Supplementary Figure 4b) (http://nevina.nve.no).

### Vegetation surveys

Vegetation surveys took place between 24^th^ July and 10^th^ of September 2020 using two methods; **1)** Vegetation plots; forming a systematic grid of 1 m^2^ quadrats covering the catchment, and **2)** a 2 m vegetation survey recording all taxa present within 2 m of the lake edge. In total 129 vegetation plots were recorded in the lake catchment (Figure 1; Supplementary Data 4).

A subset of 25-27 vegetation plots per transect were placed at varying distances from the lake shore and subsequently grouped into five distinct distance categories based on their proximity to the lake (Figure 1). Vegetation plots were aligned with sedimentary DNA sampling transects, beginning at the lake margin and continuing into the surrounding catchment (Figure 1). Plot distribution followed local topographic features and vegetation patterns, and the resulting layout was categorized into five distance groups representing increasing spatial scale from the lake (Figure 1, Supplementary Figure 5). Although a small extension of the catchment extends approximately 500 m to the northeast (Supplementary Figure 4b), we limited our vegetation plots due to the low taxonomic diversity observed beyond the last vegetation plot near the catchment’s fringes (Figure 1; Supplementary Figure 4b). All vegetation plot transects begin at the lake margin, with starting points located between 0 and 3 meters from the shoreline, depending on local topography and vegetation cover (Supplementary Figure 5). Distances from each vegetation plot to the lake shore were calculated in R by summing the distances between sequential plots along each transect (Supplementary Data 6). Based on these values, plots were classified into five distance categories reflecting increasing distance from the shoreline, with approximate ranges of less than 3 m (Distance category 1), 15-70 m (Distance category 2), 36-140 m (Distance category 3), 62-216 m (Distance category 4), and 92-284 m (Distance category 5). On average, plots within the same vegetation plot transects (Supplementary figure 5) were spaced 43.5 m apart, with distances ranging from 15.3 to approximately 76.5 m (Supplementary Data 6).

All vascular plant taxa in each vegetation plot were recorded, and taxa that were not identified in the field were collected for identification and archiving in the Tromsø University Museum herbarium (TROM). Between the two surveys, a total of 110 specimens were collected for identification and preservation. The second, more detailed, vegetation survey was within 2 m of the lakeshore (Figure 1), as the taxa closest to the lake are expected to contribute most of the eDNA ^23^. The abundance of the taxa recorded was categorized as rare **(1)**, scattered **(2)**, common **(3)** and dominant **(4)** following the definition by Alsos *et al.* 2018^23^. In addition, taxa observed outside the 2 m zone, but not observed in the vegetation plot surveys were documented, but did not receive an abundance score. The taxonomic identification for both vegetation surveys were standardized (Supplementary Data 4).

### Sediment Sampling, DNA extraction and amplification

Sediment sampling took place between 24-27^th^ of July 2020. The samples were collected in 12 northeast to southwest transects, with each sampling location 25 to 35 m apart. Two surface samples were collected per location. These were collected 15 cm apart and were recovered using a modified Kajak corer (mini gravity corer ^23^). Each surface sample was 3 cm in diameter and 2 cm thick and was subsequently stored in a 50 ml falcon tube and sample replicates from the same location were labeled as A or B. In total, 84 surface samples were collected from 42 sampling locations. The surface samples were extracted in the modern DNA laboratory at the Arctic University Museum of Norway in Tromsø. Each sample was subsampled to consist of ∼0.30 g of sediment. The subsamples were extracted using a modified DNeasy PowerSoil kit (Qiagen, Germany) protocol (QIAGEN:12855-100) following the methods described by Alsos et. al (2020) ^27^. In addition, one sediment subsampling control and seven DNA extraction controls were included.

Amplification of DNA and control extracts was performed in a dedicated ancient DNA facility to minimize contamination, using g-h primers targeting the vascular plant *trn*L (UAA) intron p6-loop locus of the chloroplast genome ^28^. The g-h primers were uniquely dual-tagged with 8 or 9 base pair tags, modified from^29^, to ensure sufficient complexity while sequencing (Supplementary Data 7). Furthermore, six PCR negative and six PCR positive controls were included for the DNA amplification. Eight PCR replicates were generated from each sediment sample or control in line with the protocol established by Voldstad et. al (2020) ^30^. After amplification, material from each sample was pooled together and cleaned ^30^. This resulted in three pools that were subsequently converted into DNA libraries using the Illumina TruSeq DNA PCR-Free protocol (Illumina Inc., CA, USA). Each of the three libraries were sequenced at ∼10% of 2 x 150-cycle mid-output flow cell on the Illumina NextSeq platform at the Genomics Support Centre Tromsø at The Arctic University of Norway.

### Bioinformatics and statistical analyses

The raw sequence data was analyzed with the OBITools software package^31^ as described in the methods by Rijal et al. (2021)^32^. We identified the barcodes with four reference databases: The global EMBL database (release 143), PhyloAlps^33^, PhyloNorway ^5,34^, and a local taxonomic reference database, ArcBorBryo, containing boreal and arctic vascular plant taxa and arctic bryophytes^35–37^.

We removed sequences from our data if they were identified as PCR artifacts by *obiclean*, with less than three reads per observation or less than 10 reads across the entire dataset. Sequences were removed if they appeared more frequently in the negative controls compared to the samples. Sequences assigned to the same taxon were collapsed by combining their read counts and retaining the maximum number of PCR repeats. For the identification of the barcodes, we prioritized the databases in the following order: PhyloNorway, ArcBorBryo, PhyloAlps and EMBL. Uncertain identifications were manually inspected by blasting the sequence to the NCBI nucleotide database. The final taxonomic assignment of the retained taxa were categorized as 1 (assumed true positive as expected in the region and/or recorded in the vegetation surveys), 2 (likely true positive as biogeographically marginal and/or sequence variant of more abundant sequence), 3 (assumed either false positive or food contaminant). Only category 1 and 2 taxa were used for further analysis (Supplementary Data 1).

The analytical quality scores ^32^ were used to identify samples that had poor replicability for the identified taxa. The similarity between the A and B samples was investigated by calculating the Jaccard index using the presence-absence taxonomic assignments rather than the barcodes. Then for subsequent analysis, the A and B samples for each sampling location were collapsed into one sample and any taxon that was not replicated was removed (Supplementary Data 3). For each taxon, the frequency of detection was calculated for the 42 lake sampling locations.

To get an estimate of the optimal number of samples required to capture optimal diversity information from the lake, taxa accumulation curves were generated with 1000 randomizations for the 42 merged DNA samples. A breakpoint analysis was performed on the accumulation curve to estimate an optimal number of samples to maximize plant diversity capture of the catchment. We fitted a linear regression model with accumulated species number as the response and the number of samples as the predictor which was then subjected to segment analysis to find breakpoint in the species accumulation using the segmented package ^38^ in R (R Core Team 2024; Figure 2a).

Both DNA and vegetation lists were standardized to compare the sedDNA results with the vegetation taxa list. When taxa were detected at different taxonomic levels between the two datasets, for example due to the taxonomic resolution of the marker used, the identification reverted to the highest common taxonomic level. The frequency of the taxa in the sedDNA results were compared to both the 2 m vegetation surveys and the vegetation plots. The frequency of taxa in sedDNA and vegetation plot survey were calculated as the sum of the detections of each taxon across 42 merged DNA samples and across 24 vegetation plots (after merging variants of plot no 13, see Supplementary Data 4) within each distance category respectively. We considered the abundance estimate from the 2 m vegetation survey and taxon frequency from vegetation plots as two independent proxies for biomass.

The association between the sedDNA and the vegetation data was evaluated using linear models. We obtained a reasonable model fit when the DNA frequency was considered as the response and vegetation abundance from the 2 m survey as the categorical predictor variable (Supplementary Figure 6). The DNA frequency was square root transformed when vegetation frequency from Distance categories, 1 (less than 3 m from lakeshore), 4 (approximately 62-216 m from lakeshore), and 5 (approximately 92-284 m from lakeshore) were considered as the predictor. Both the response and predictor variables were log transformed (variable+1 to avoid producing infinite values) while assessing the associations between DNA detection and vegetation frequencies at Distance categories 2 (approximately 15-70 m from lakeshore) and 3 (approximately 36-140 m from lakeshore). Both the response and predictor variables were square root transformed while assessing the associations between DNA detection and combined vegetation frequencies from all distance zones (Supplementary Table 2, Supplementary Figure 6). All the analyses were performed in R package^38^, model assumptions were checked using DHARMa package ^39^, and results were visualized using ggplot package ^40^.

### Data Accessibility

The raw DNA sequence data generated have been deposited in the European Nucleotide Archive (ENA) under BioProject accession code PRJEBXXXXXX. The identified OBITools output is available on Dryad: DOI:XXXXX.

### Code availability

The R scripts used for statistical analyses and figure generation will be made available upon acceptance at a public GitHub repository and archived with a DOI via Zenodo.

## Acknowledgements

We thank Galina Gusarova for her assistance with the identification of the collected vegetation specimens. Amy McDermott and Scarlett Zetter for their aid in collecting the sediment samples. Marie Føreid Merkel for assisting with the lab work. Luke Dane Elliott for analyzing the metabarcode data. Tulug Gulce Ataman was funded by an internal PhD position at The Arctic University Museum of Norway. Bioinformatic analyses were performed on resources provided by UNINETT Sigma2 - the National Infrastructure for High-Performance Computing and Data Storage in Norway.

## Author contributions

T.G.A and A.G.B conceptualized the study. T.G.A. designed the experimental plan and generated the data. T.G.A., Y.L., I.G.A. and D.P.R. performed the analysis of the data. T.G.A. wrote the manuscript with feedback from all co-authors.

