## Supplementary Materials for "Maximizing vegetation representation of the catchment in sedimentary DNA with targeted cores in small lakes"

#### 1. Jaccard similarity distribution

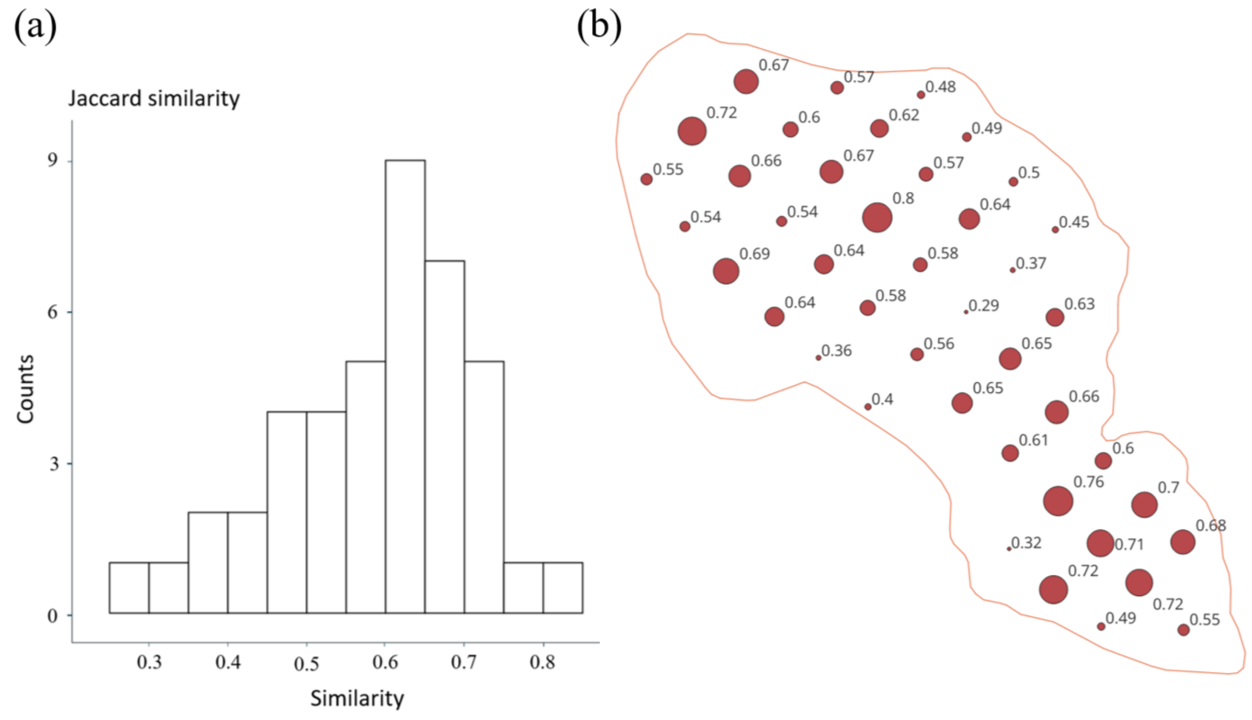


**Supplementary Figure 1: (a)** A similarity histogram based on Jaccard similarity values obtained from the A and B sample comparisons. x-axis shows the percentage of similarity, y-axis shows the number of sample pairs. **(b)** Jaccard similarity distribution based on merged pairs (shared taxa/Total richness) Sample points are proportional to Jaccard similarity levels in each 42 sampling pairs demonstrated in the map.

### 2. Correlation between aquatic reads vs richness and similarity

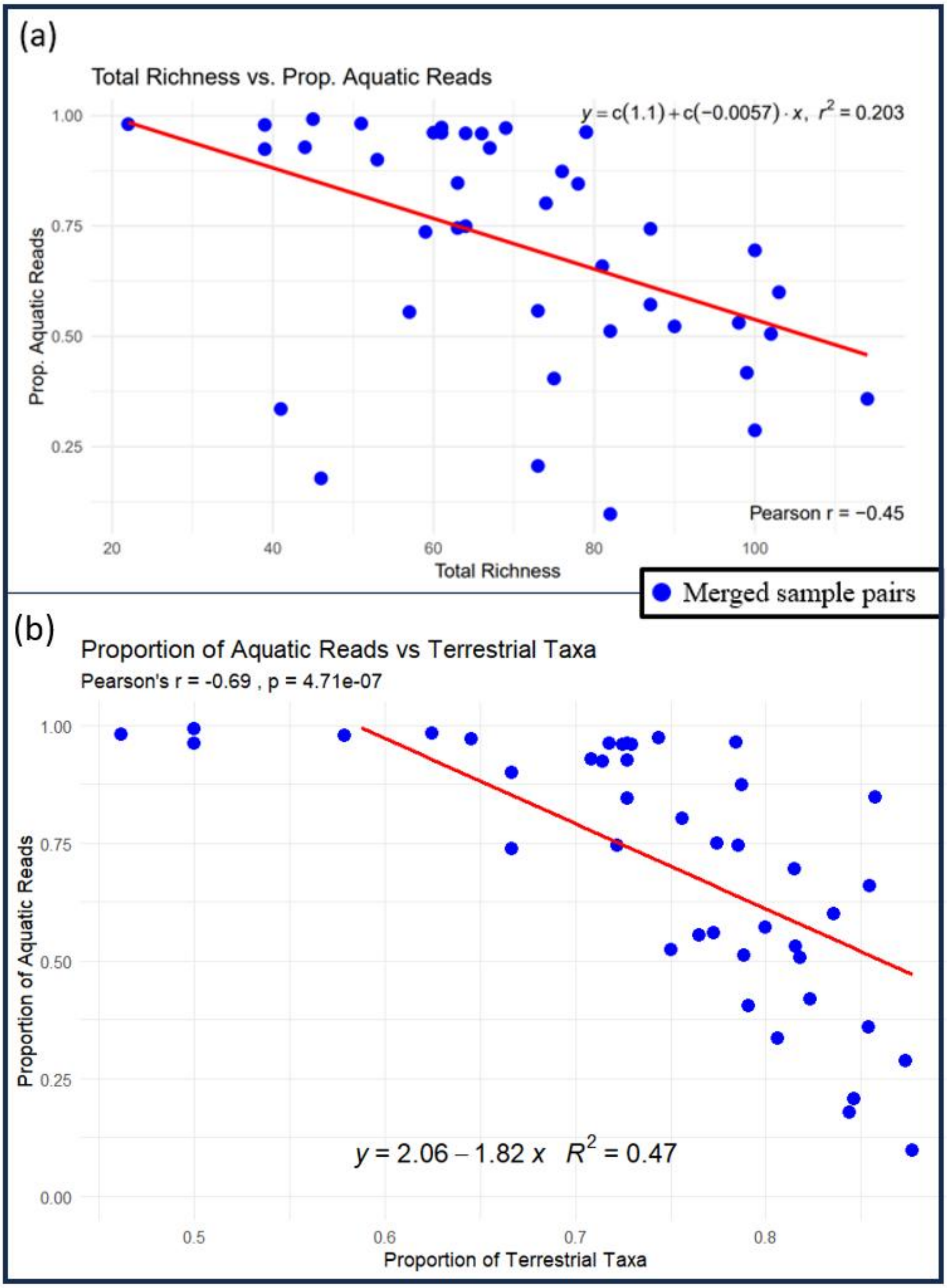


**Supplementary Figure 2:** Pearson correlation between proportion of aquatic reads in merged A and B samples (a) Correlation between total richness (terrestrial and aquatic taxa in merged samples) vs aquatic reads from 42 merged samples (b) Proportion of terrestrial richness (shared terrestrial/shared(A∩B) as shared richness in merged samples) vs proportion of aquatic reads

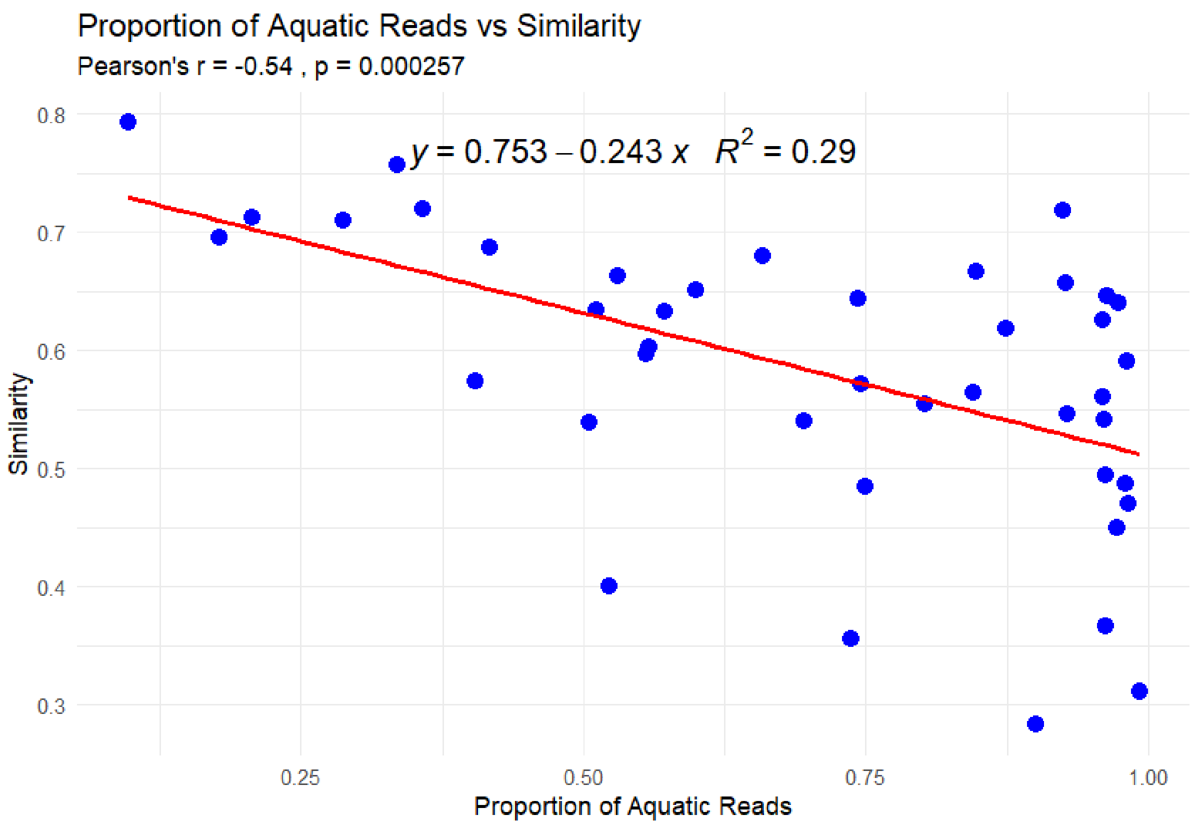


**Supplementary Figure 3:** Relationship between replicate similarity and the proportion of aquatic reads in sediment samples. Each point represents a merged sample pair, with the proportion of aquatic reads on the x-axis and replicate similarity on the y-axis. The red line shows the linear regression fit ($y = -0.24x + 0.75$, $R^2 = 0.29$), indicates a significant negative correlation (Pearson's $r = -0.54$, $p = 0.00026$). Samples with more aquatic reads tend to show lower replicate similarity.

### 3. Catchment Delineation and Comparison

Catchment boundaries were delineated using the NEVINA platform (https://nevina.nve.no), which generates watersheds from a national 10 m Digital Elevation Model (DEM) based on a user-defined pour point. Because NEVINA is sensitive to the pour point location, multiple boundaries were tested. Outputs varied notably: some included only one side of the lake or extended well beyond topographic ridgelines. It was noticed that the hydrological catchment did not follow the topography due to extensive land-drainage. One such configuration produced a catchment of 0.8 km², which we used as a reference in our comparisons (Supplementary Figure 1a). Therefore, we re-delineated manually based on field observations of the drainage pattern and flow directions. This re-delineated catchment, is shown in Supplementary Figure 1b (dashed line), covers 0.6 km². The grid overlay highlights the northern and northeastern areas where vegetation was sparse and largely composed of grasses. All vegetation survey plots were located within this manually defined catchment. We extracted standard field parameters from NEVINA for both catchments to assess key differences. These include land cover proportions, slope, drainage density, field length, hypsographic elevation distribution, and modeled climatic values such as average annual runoff (1991–2020). The smaller, custom catchment includes a higher percentage of cultivated land (41.6% vs. 22%), a shorter field length (0.7 km vs. 1.4 km), and slightly steeper average slope (8.7° vs. 8.3°). Both catchments show similar elevation ranges, drainage densities, and runoff values.

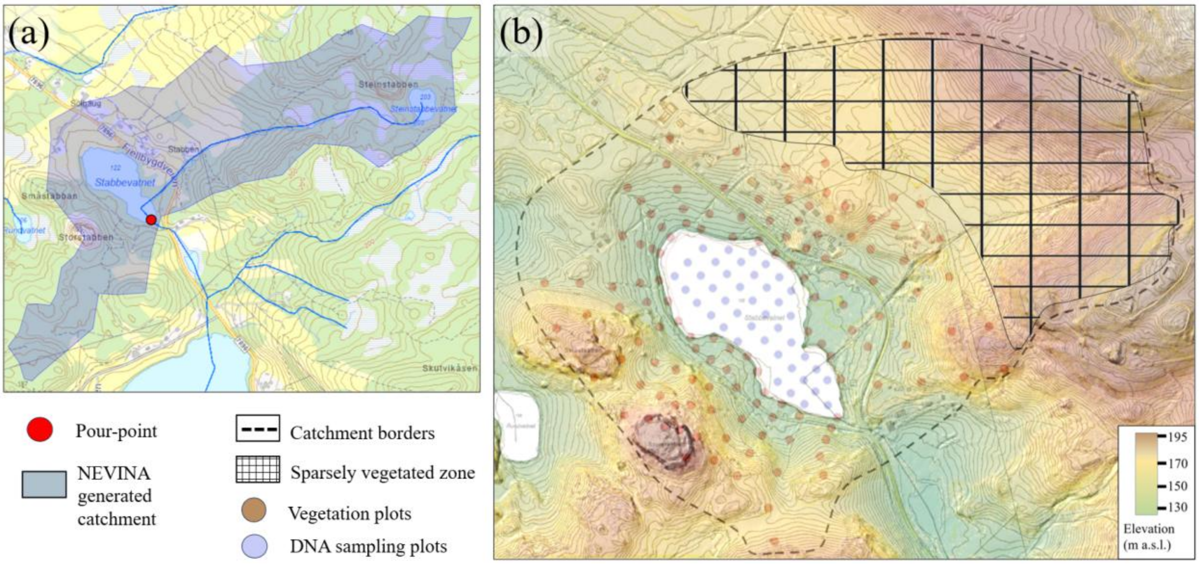


**Supplementary Figure 4: Catchment Delineation and Comparison (a)** A configuration of the catchment area generated using NEVINA (blue shading), based on a selected pour point (red), without manual modification. **(b)** Catchment boundary (dashed line) manually delineated based on field observations and topographic context, representing the area most likely to contribute terrestrial material to the lake. Vegetation plots are located within this boundary, extending from the northeastern edge to the southern shore. The northern and northeastern sections included sparse vegetation (grids), mainly grass, and contributed little to the overall vegetation signal. Elevation contour lines were shown at 10 m intervals.

### 4. Vegetation plot transects

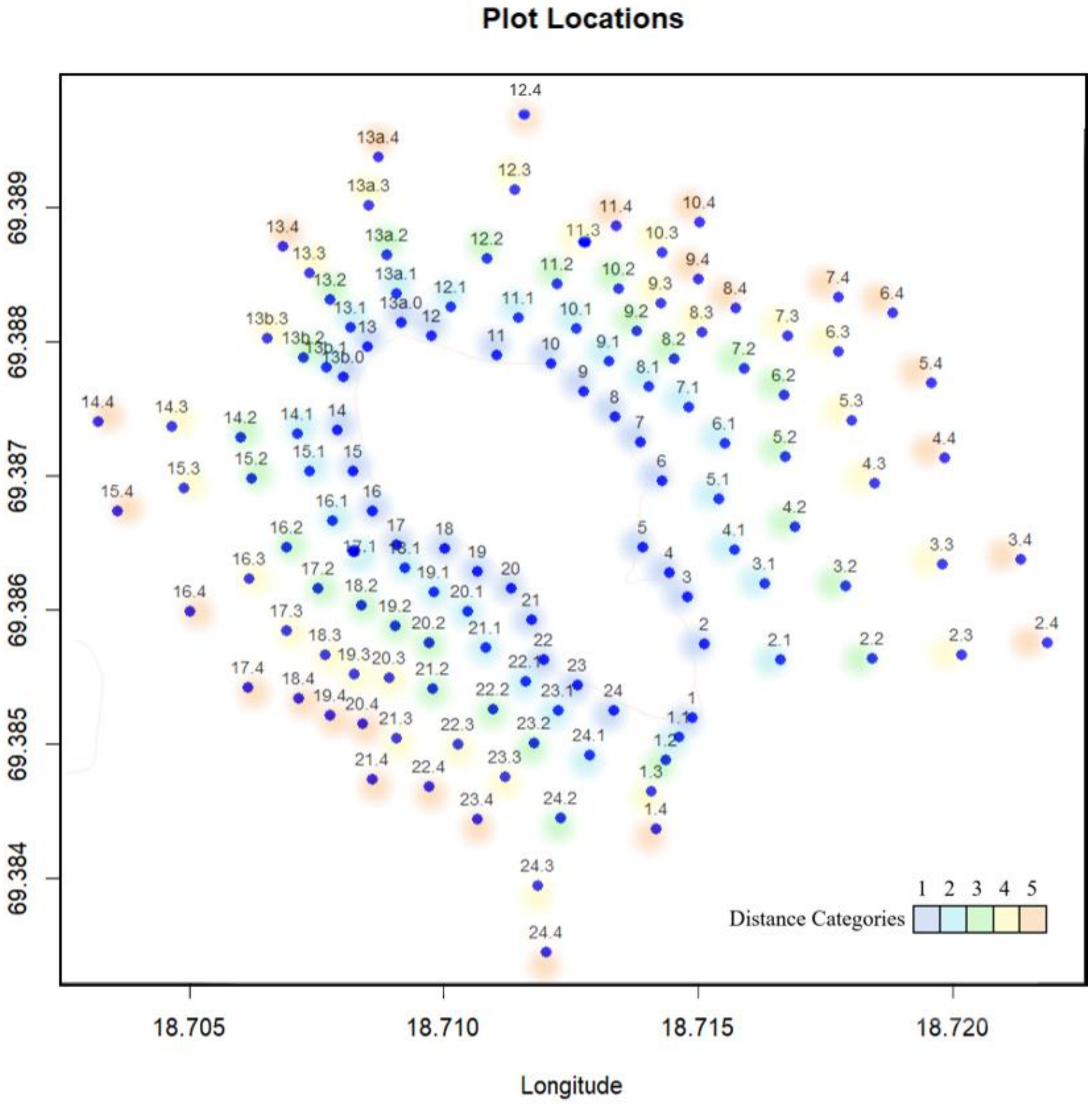


**Supplementary Figure 5. Map of vegetation plot locations**. The x-axis shows longitude and the y-axis shows latitude. Vegetation plots are organized in transects (e.g., 1.0 to 1.4), each beginning near the lake margin (0–3 m from shore) and extending inland across five distance zones. Plots (blue) were positioned along straight lines parallel to the sedimentary DNA sampling transects, with spacing and endpoints determined by topography.

### 5. Diagnostic plots for regression models

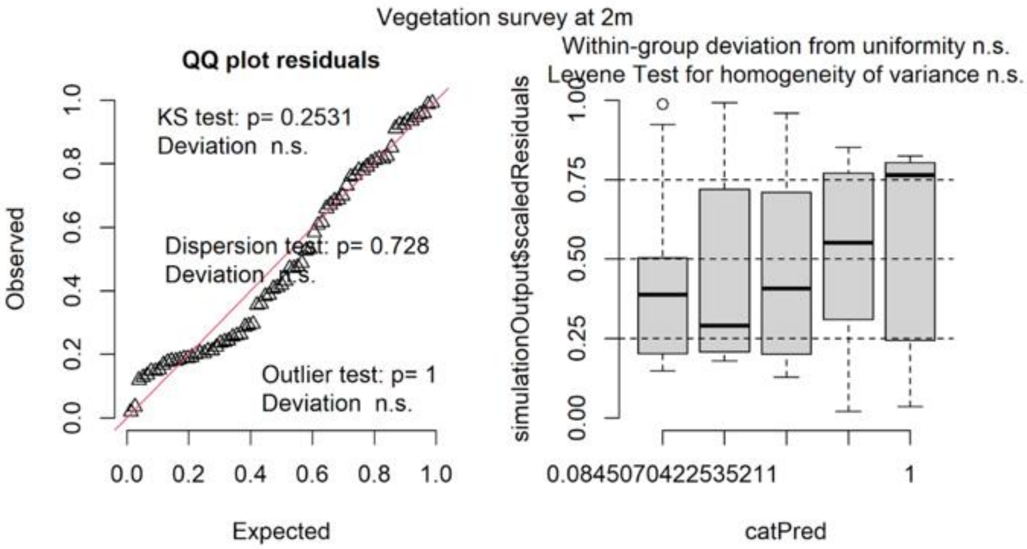


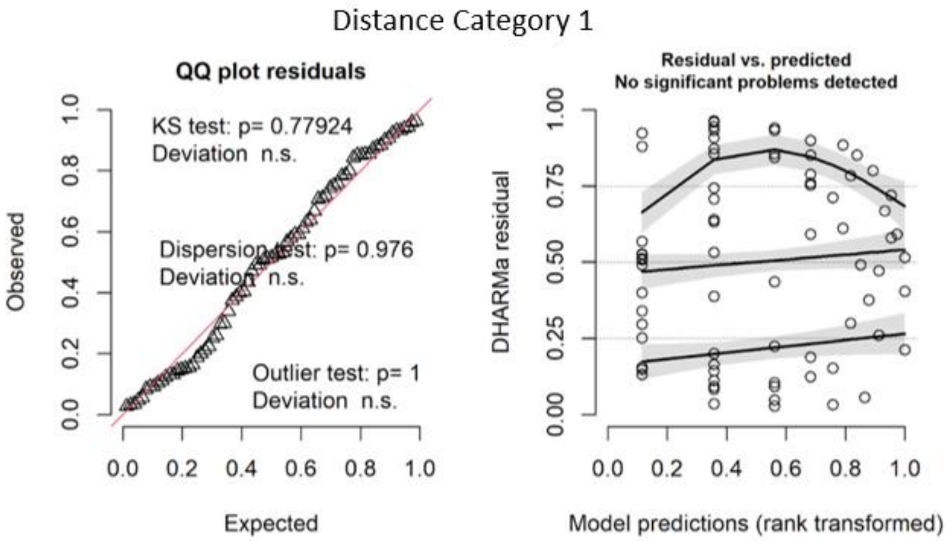


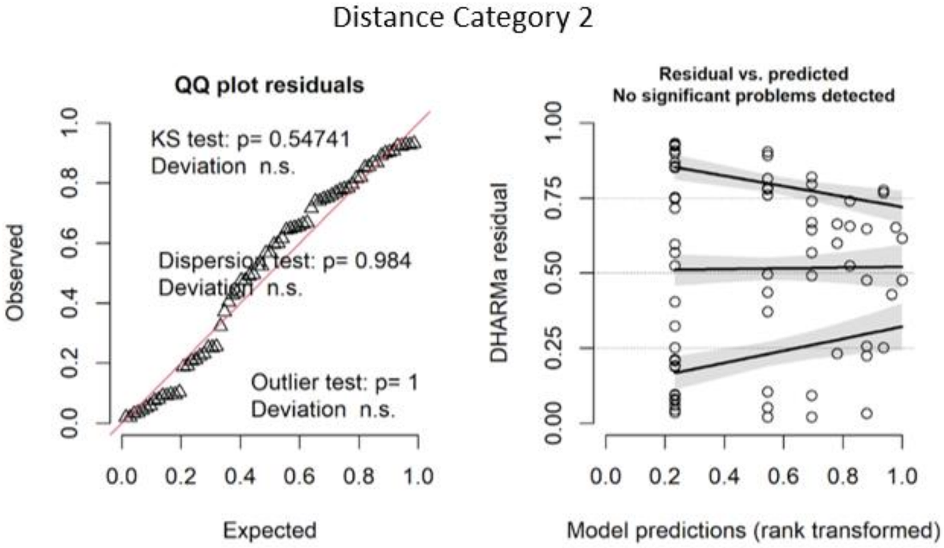

### Distance Category 3

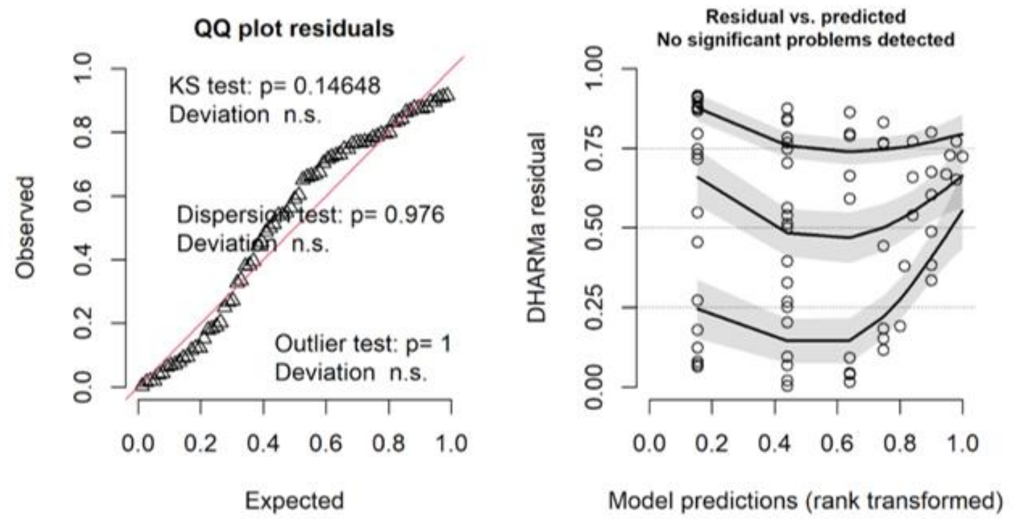


### Distance Category 4

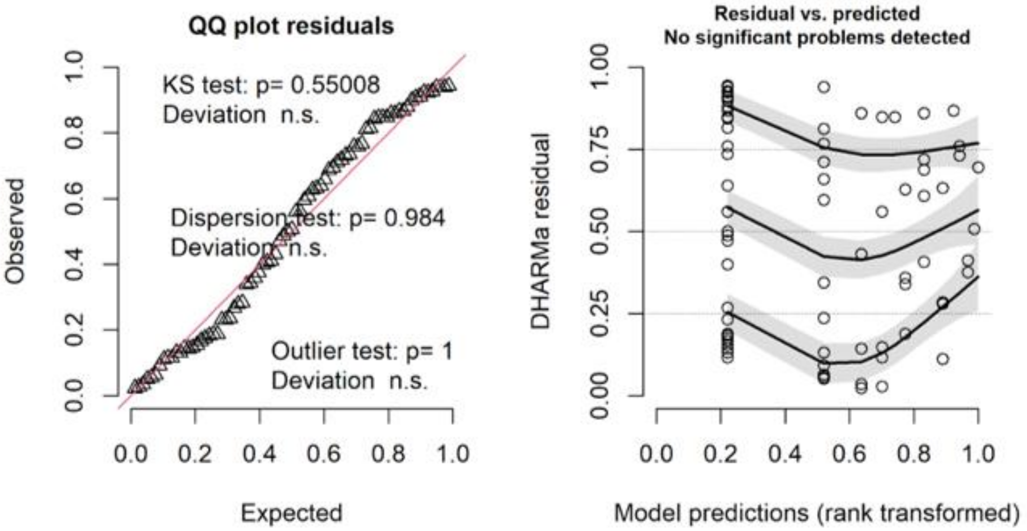


### Distance Category 5

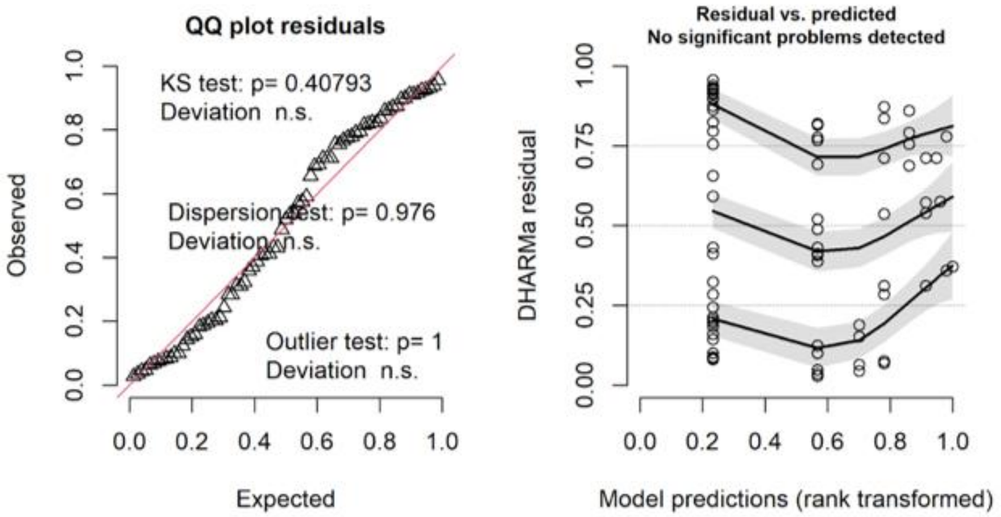

**Supplementary Figure 6:** Diagnostic plots for regression models. DNA frequency was treated as the response and ranked abundance from the 2 m vegetation survey as well as vegetation frequency from plots of different distances were treated as the predictor variables.

### 6. Summary Statistics

| Abundance category | Parameter | Estimate | Std..Error | t.value | P-value |
|---|---|---|---|---|---|
| 0 (absent) | Intercept | 11.88 | 3.04 | 3.91 | **0.000** |
| 1 (rare) | Slope | -0.88 | 4.76 | -0.18 | 0.855 |
| 2 (scattered) | Slope | 1.08 | 3.96 | 0.27 | 0.785 |
| 3 (common) | Slope | 17.88 | 4.30 | 4.16 | **0.000** |
| 4 (dominant) | Slope | 19.57 | 5.07 | 3.86 | **0.000** |

**Supplementary Table 1:** Summary statistics from a linear regression models where DNA frequency was treated as the response and vegetation abundance from 2m vegetation survey as the categorical predictor. All the p-values $< 0.05$ at 4 and 70 degrees of freedom are indicated in bold.

| Distance Category | Transformation | Parameter | Estimate | Std.. Error | t.value | p-value | R_square | F_val | Degrees of freedom |
|---|---|---|---|---|---|---|---|---|---|
| Distance Category 1 | Square root | Intercept | 3.15 | 0.27 | 11.81 | 0.000 | | | |
| Distance Category 1 | Square root | Slope | 0.16 | 0.04 | 3.89 | **0.000** | 0.17 | 15.102 | 1& 73 |
| Distance Category 2 | Log-Log | Intercept | 2.34 | 0.16 | 14.62 | 0.000 | | | |
| Distance Category 2 | Log-Log | Slope | 0.37 | 0.15 | 2.49 | **0.015** | 0.08 | 6.212274 | 1 & 69 |
| Distance Category 3 | Log-Log | Intercept | 2.26 | 0.20 | 11.17 | 0.000 | | | |
| Distance Category 3 | Log-Log | Slope | 0.24 | 0.16 | 1.44 | 0.155 | 0.03 | 2.068979 | 1& 73 |
| Distance Category 4 | Square root | Intercept | 3.37 | 0.29 | 11.75 | 0.000 | | | |
| Distance Category 4 | Square root | Slope | 0.11 | 0.06 | 1.78 | 0.079 | 0.04 | 3.162712 | 1 & 75 |
| Distance Category 5 | Square root | Intercept | 3.54 | 0.28 | 12.66 | 0.000 | | | |
| Distance Category 5 | Square root | Slope | 0.13 | 0.09 | 1.46 | 0.149 | 0.03 | 2.125139 | 1& 73 |
| Distance Category 12 | Log-Log | Intercept | 1.72 | 0.22 | 7.77 | 0.000 | | | |
| Distance Category 12 | Log-Log | Slope | 0.53 | 0.13 | 4.10 | **0.000** | 0.19 | 16.81706 | 1& 73 |
| Distance Category 123 | Log-Log | Intercept | 1.31 | 0.25 | 5.17 | 0.000 | | | |
| Distance Category 123 | Log-Log | Slope | 0.61 | 0.13 | 4.72 | **0.000** | 0.22 | 22.28614 | 1 & 77 |
| Distance Category 1234 | Square root-Square root | Intercept | 1.73 | 0.40 | 4.28 | 0.000 | | | |
| Distance Category 1234 | Square root-Square root | Slope | 0.63 | 0.13 | 4.89 | **0.000** | 0.23 | 23.90647 | 1 & 81 |
| Distance Category 12345 | Square root-Square root | Intercept | 1.56 | 0.40 | 3.91 | 0.000 | | | |
| Distance Category 12345 | Square root-Square root | Slope | 0.62 | 0.12 | 5.15 | **0.000** | 0.24 | 26.47814 | 1 & 84 |

76

**Supplementary Table 2:** Summary statistics from linear regression models where DNA frequency was treated as the response and vegetation frequencies from plots of distinct distant Distance Categorys were considered as predictors. All the statistically significant ($p < 0.05$) associations are indicated in bold.
